# Adaptive dynamics of unstable cancer populations: the canonical equation

**DOI:** 10.1101/217851

**Authors:** Guim Aguadé-Gorgorió, Ricard Solé

**Affiliations:** ICREA-Complex Systems Lab, Universitat Pompeu Fabra, 08003 Barcelona, Spain; Instituí de Biología Evolutiva (CSIC-UPF), Psg Maritim Barceloneta, 37, 08003 Barcelona, Spain and; Santa Fe Institute, 399 Hyde Park Road, Santa Fe NM 87501, USA

**Keywords:** Cancer evolution, genetic instability, Adaptive Dynamics, canonical equation

## Abstract

In most instances of tumour development, genetic instability plays a role in allowing cancer cell populations to respond to selection barriers, such as physical constraints or immune responses, and rapidly adapt to an always changing environment. Modelling instability is a nontrivial task, since by definition evolving changing instability leads to changes in the underlying landscape. In this paper we explore mathematically a simple version of unstable tumor progression using the formalism of *Adaptive Dynamics* (AD) where selection and mutation are explicitly coupled. Using a set of basic fitness landscapes, the so called *canonical equation* for the evolution of genetic instability on a minimal scenario associated to a population of unstable cells is derived. The implications and potential extensions of this model are discussed.

## I. INTRODUCTION

Cancer can be understood as the failure of those regulatory mechanisms that guarantee the maintenance of tissue and organ homeostasis. Cooperative interactions, along with extensive feedback signalling loops and replication checkpoints provide multiple paths to avoid the emergence of undesirable mutations or chromosomal abnormalities that can allow rogue cells to start proliferative growth. In dynamical terms, what has to be avoided within multicellular organisms is any kind of individual cell Darwinian evolution (Nowell, 1976; Greaves et al., 2012; Gatenby et al., 2017).

It is generally acknowledged that genetic instability plays a key role in tumour progression and carcinogenesis (Hanahan et al., 2011). Unstable genomes result from the failure of molecular mechanisms responsible for the maintenance of genome integrity (Negrini et al., 2010). That cancer cells are unstable is fairly well illustrated by the observation of their karyotypes: in sharp contrast with healthy cells, cancer chromosomal arrangements reveal a wide degree of aneuploidy (Lengauer et al., 1998). Such high levels of mutational load deploy the potential to overcome selection barriers, as well as involve a rather uncommon process from multicellularity to reduced cellular complexity (Solé et al., 2014), giving place to a highly adaptive and heterogeneous population. Genetic instability acts as a driver as well as the search engine for disease progression. An important (and not always appreciated) consequence of instability is that, once unleashed, can easily grow as the lack of proper repair can damage other components of the check-and-repair cellular network.

The fact that genetic instability itself changes over cancer evolution makes it difficult to properly model its behaviour. A first approximation is to reduce instability to a rate, which is assumed to be fixed. By tuning it through its range of possible values, different phases and transitions appear and can be analysed. However, a changing instability rate affects, by means of modifying the probability of mutations, all kinds of replication and control mechanisms within the vast pathways towards cancer malignancy. Within this picture, instability cannot be taken as a parameter, but rather as an evolving phenotypic trait affected by the selective pressures of the tumor microenvironment.

Cancer has a complex evolutionary nature (Merlo et al., 2006), with adaptive medicine being a promising step towards novel cancer treatments (Gatenby et al., 2009; Gillies et al., 2012; Ibrahim-Hashim et al., 2017). Evolutionary models of cancer have been developed but only small steps have been made on the evolutionary dynamics of genetic instability (Datta et al., 2013; Asatryan et al., 2016), a hallmark of cancer with a complex molecular basis (Negrini et al., 2010; Loeb, 2001). We propose that the evolutionary paths followed by unstable populations are describable by means of the framework of Adaptive Dynamics (AD) (Dieckmann et al., 1996; Champagnat et al., 2001) which has been used so far in the study of cancer niche construction (Gerlee et al., 2015). AD models provide a powerful alternative to previous formal approaches by explicitly including replication, mutation and selection in a consistent way, allowing an exploration of the evolutionary dynamics of adaptive traits.

A central object in the AD framework is the so called *canonical equation.* For a given quantitative phenotypic trait s, this equation describes the evolutionary trajectory for the mean trait value as

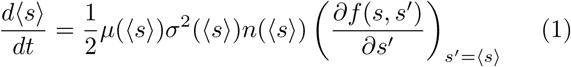

where µ*(〈S〉)* is the probability that a mutant individual (for the given trait) will be generated, σ^2^ is the variance of the distribution for mutants *s′* derived from an individual with trait *s*, *n* the stationary population size and the last term in the rhs stands for the fitness gradient associated to the specific landscape at work. The standard formulation involves some given assumptions on the mutation-selection process, and we will therefore review the mathematical process in order to understand up to which point the framework is suitable for our problem. Understanding how instability changes can give further insight in understanding its role as a cancer hallmark, and might as well produce relevant steps towards contemplating genetic instability as potential target for treatment. Is it possible to formulate a canonical equation describing the time evolution of instability? The answer is affirmative and here we show how it can be obtained.

## II. POPULATION DYNAMICS

With the aim of obtaining a clear understanding of the questions proposed above, we look for a minimal model to implement the unstable evolutionary dynamics. Our goal is to consider the process of cancer progression, which involves a heterogeneous population of cells (figure 1a). Since we look for a minimal set of conditions, a so called Moran process (Moran, 1958) will be considered. Here the total population of cells will be assumed to remain constant in time, while replication, death and mutation processes occur. The starting point to build our model is a set of differential equations describing a population of replicators, namely:

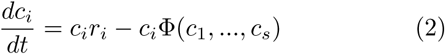

where *c_1_,…c_s_* indicates the different population abundances for each strain (a finite set of strains is assumed). Each population replicates with a rate *r_i_* and Φ(c_1_,…, c_s_) is a competition term that incorporates the presence of growth constraints. In order to implement the constant population constraint imposed by the Moran process, we need to impose Σ_*j*_ *c*_*j*_ (*t*) = *N* where *N* stands for the total population size. Since

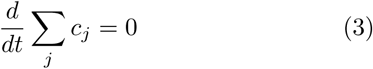

this condition implies that this imposes 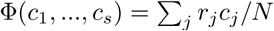 and thus the competition model

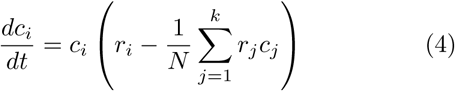

represents the mean-field, well mixed competition model.

**FIG. 1:**
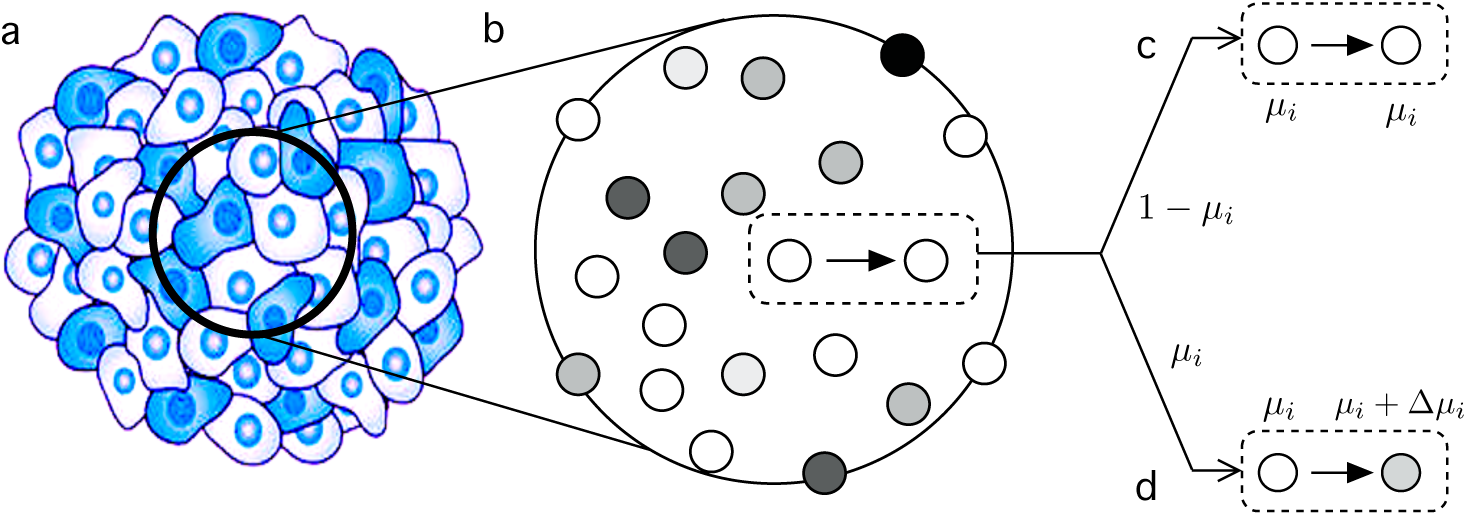
The Moran process rules associated to the model of a population of unstable cells competing for resources. We consider an idealised model of a heterogeneous cancer cell population (a) described by a well-mixed (mean field) model (b). Here cells occupy a given domain Ωthat is not explicit and each cell has a distinct phenotype described in particular by its intrinsic instability µ_*i.*_ In the Moran process, when a cell replicates it occupies another cell’s niche and produces an identical daughter (c) or a slightly different one due to a mutation event proportional to *yi,* which can lead to an increase of the instability levels (d).

A particularity of the Moran process is that a cell of type *ci*gives birth by means of occupying another cell’s site with probability *r_i_*, so that the birth-death process is coupled into a single event that will eventually lead to selection towards cells with higher *r_i_*. Furthermore, mutation is introduced by considering that cells can give birth to mutant offspring with probability *μ_i_.*

Mutations, however, do not occur as in quasispecies or replicator-mutator models, where genomes mutate from one to another. In our model, a newly-born mutant cell will have a modified mutation rate *μ′ = μ_i_ + Δμ,* where Δ*μ* is taken from a continuous distribution that we discuss later on. With this, we emphasize the wide levels of heterogeneity and genomic configurations found within tumours by means of giving a diferent phenotype to each cell rather than grouping populations into a countable, finite set of possible genome configurations.In light of this, a mean-field approach is no longer useful to treat this model.

Our goal is to understand how are selection and mutation coupled when instability, and thus the individual mutation rate *μ_i_*, can itself change and affect the rate of cell replication *r_i_*, and what are the evolutionary consequences of this coupling.

## III. SELECTION ON INSTABILITY

Multiple molecular mechanisms affect the genome stability of a cell, eventually leading to variations in the fidelity of DNA replication (Vogelstein et al., 2002). Here we simply assume that, when a cell reproduces (with probability *r*_*i*_) and undergoes a mutation (with probability μ_*i*_)), the offspring cell can experience a change in the mutation rate itself, due to, for example, mutations having occured on a gene related to DNA stability (Lengauer et al., 1998) or oncogene mutations inducing further replicative stress and DNA damage (Negrini et al., 2010).

Moreover, it is accepted that genome instability is a hallmark of cancer (Hanahan et al., 2011) since its role as the driver towards the alterations that result in tumor malignancy. A most common event during this process are mutations in oncogenes that usually result in increased levels of replication (Vogelstein et al., 2002). On the other hand, elevated levels of instability can trigger deleterious mutations in house-keeping genes leading to reduced cell viability or death. A coupling exists between replicative capacity, cell viability and mutation rate. We propose a minimal adaptive landscape that translates such coupling into replication rates being a function of instability, r(µ).

### A. Adaptive Landscape

Within our minimal scope we consider that mutations on oncogenes can translate into a linear increase in replication rythmn, such that r(µ) = r_0_ *+ N_R_δ_R_µ,* with r_0_being the basal replication rate of normal cells, *N_R_* the number of oncogenes responsible for increased replication and *δ_R_* the mean effect on replication rate when mutating one of such genes. In this picture, however, we need to take into account the minimal genetic material needed for a cell to keep its basic functions. If we group such material into a number of house-keeping genes, the probability that none of them has been mutated is (1-µ)^NHK^. Grouping both considerations together we obtain an analytical description of the coupling between replication and instability

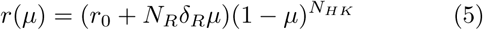

(Solé et al., 2014). This adaptive landscape is of course of qualitative nature, and realistic fitness landscapes for unstable tumor environments are still far from our knowledge. However, certain points can be made if we give values within realistic parameter ranges to our function. The number of both oncogenes and housekeeping genes have been widely assessed, and we take them to be about N_R_ ≈ 140 (Vogelstein et al., 2013) and *N_HK_ ≈* 3804 (Eisenberg et al., 2013) respectively. Interestingly enough, considering small replication effects for *δ*_R_, such experimental values produce an adaptive landscape (Fig. 2) that has a positive gradient within the region of µ, **G** [10^-9^,10^-4^], so that our evolutionary trajectories will be bounded within a region of mutation rate values in accordance with those experimentally measured for tumor cells (Tomlinson et al., 1996).

**FIG. 2:**
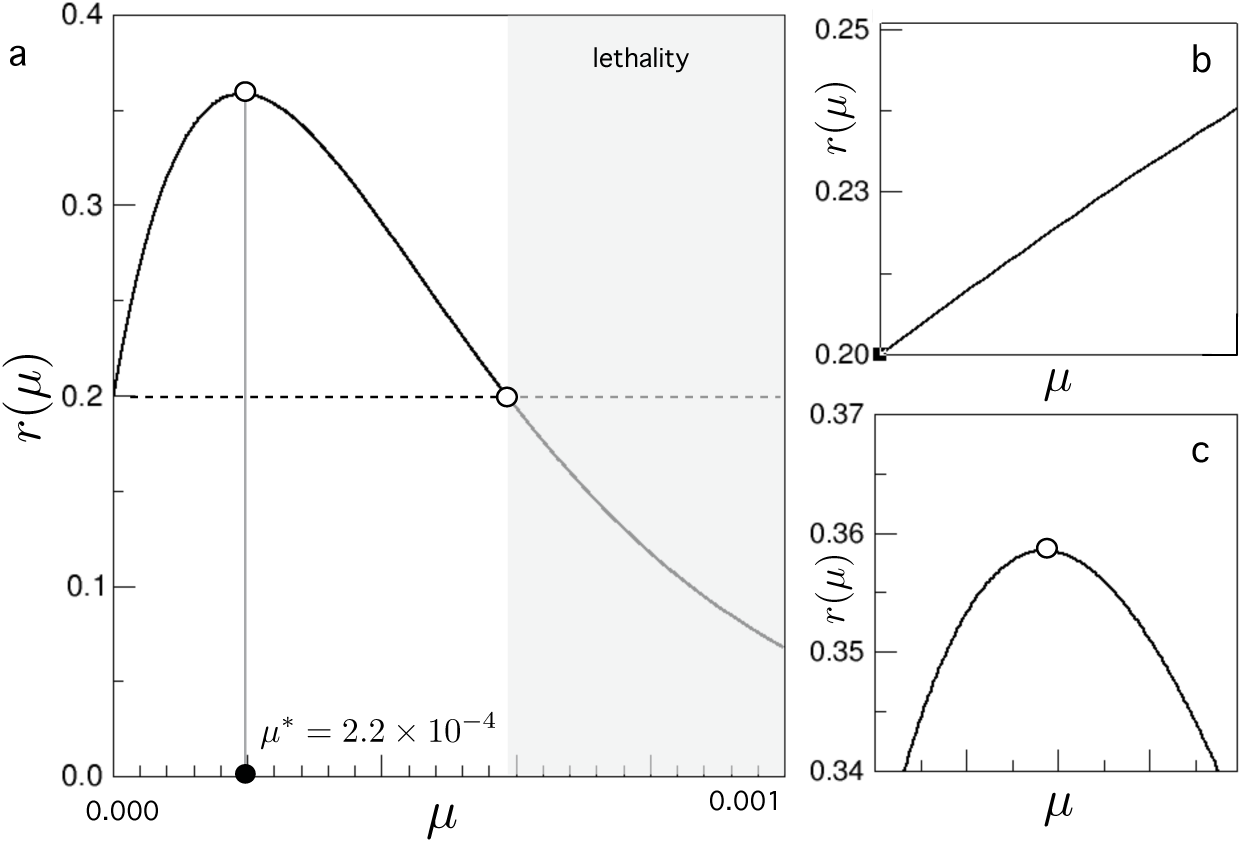
Fitness landscape function associated to the Moran process of unstable tumour cells. In (a) the full landscape, given by a replication rate *r(μ)* = *(r0 + N_R_S_R_p)(l — P)^NHK^*, is plotted against the instability rate *μ.* Further discussion is focused on two limit cases representing initial linear progression of instability (b) and optimal mutation (c) domains.

### B. Distribution of new mutations

We have assessed so far what is the effect of instability in proliferation, thus coupling mutation and selection for mutation rate. Up next, we need to evaluate how does instability change during reproduction, so that we can finally compute the effects on replicative capacity of a mutated cell. As previously discussed, a broad range of mechanisms relate to variations in DNA replication fidelity. Such variations, however, are hardly in the direction of increasing DNA stability, and in general account for an increase in the mutation rate of cancer cells due to accumulation of further tumor-supressor or care-taker gene mutations. (Vogelstein et al., 2002)

This trend of generating more unstable offspring is translated into a positively skewed distribution of mutants *M*(µ, Δµ). To keep the mathematical background of our model treatable, a Rayleigh distribution peaked at Δμ, = 0 has been chosen ^1^. Under this scheme, instability of a daughter cell is likely to be similar or slightly higher from its parent, controled by a scale parameter 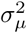 depicting the general size of mutational increases.

## IV. ADAPTIVE DYNAMICS

Adaptive Dynamics is a set of techniques or a mathematical framework that models long term phenotypic evolution of populations. Several works by different authors cover a broad scope of possible applications, and we hereby focus on the work of Dieckmann and Law and others (Dieckmann et al., 1996; Champagnat et al., 2001) and adapt it to our particular system. The main biological background behind the maths sits in considering the evolutionary step as a mutant appearing and invading in a population in ecological or dynamical equilibrium (Dieckmann et al., 1996). Under this picture, the ecological and evolutionary time scales are considered to be uncoupled, so that the process of the mutant competing against the resident population, and eventually fixating in it, is considered instantaneous in the evolutionary process.

General AD literature (see e.g. Dieckmann et al., 1996; Geritz et al., 1998; Champagnat et al., 2001) follows the evolution of a quantitative phenotypic trait or set of traits, s, that can change through mutations. In such original works, the rate _μ_ at which mutations appear is considered a possible function of the trait s, but afterwards and further on in the AD literature is usually left as a constant of each model. In the light of what we have discussed in the previous section, however, instability itself is a quantitative trait if computed as a mutation rate, and so the coupling of mutation and selection results in s = _μ_ being the studied trait value.

The starting point of the AD modeling is to consider the evolutionary process, where the population’s mutation rate changes as mutants appear and fixate, as a Markov chain for the probability of finding the population at time t having trait value _μ_

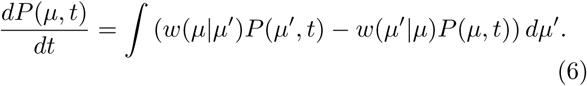

The transition probabilities *w(μ′|μ)* describe the evolutionary step and contain the probability of the mutant with trait µ*′* appearing (*A*) and fixating (ρ) in the population, so that *w(µ′*|µ) *= A(µ, µ′)ρ(µ, µ′).* The probability that a mutant appears is A(µ, µ*′) = Nr(µ)μM(µ, µ′*), the size of the population at equilibrium *N*, the probability of birth and mutation *r(μ)μ* and the probability that the mutant has mutation rate p*′* provided the parent cell had rate *μ*.

The probability *ρ(µ, µ′)* that a mutant with fitness advantage *r(µ′)/r(µ)* fixates in a population of *N* individuals has an analytical expression for the Moran model (Ewens, 2004)

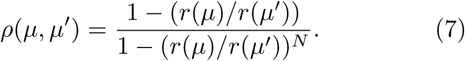

A common procedure of the AD framework is to ex-pand p for small variations of the trait value under the assumption of large populations, assumptions that are not a restriction for our problem. Under this view, the probability that the µ*′* mutant fixates is zero for r(µ*′*) ≤ r(μ) and

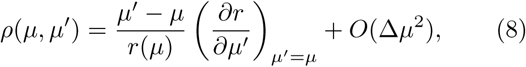

for r(μ′) > *r(μ).* Once a complete expression for the transition probabilities is build, we only need to recall how the evolution of the mean mutation rate can be written as

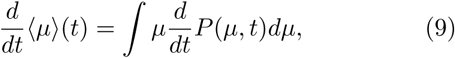

so that, using the original master equation, we obtain

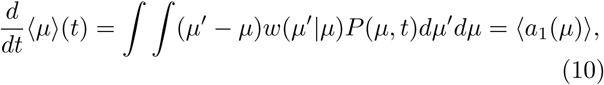

where one recalls that 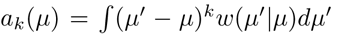 is the *k*-th jump moment. If the first jump moment were a linear function of μ, then 〈a_1_(µ)〉 = a_1_(〈μ〉), a condition that settles a well-studied range of validity even under nonlinear evolutionary dynamics (see e.g. van Kampen 1962, Kubo 1973), and will still be a good approximation within regions of small mutation rate and fitness effects. The evolutionary trajectory for the mean path (we cease denoting it by angle brackets) will therefore follow

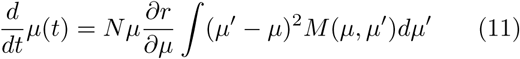

At this point, we recall that only fitter mutants can invade and so may eventually contribute to the exploration of the adaptive landscape. This translates onto the domain of the integration being restricted to μ′ *>* µ, and so we integrate the positive part of our 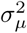 skewed mutant distribution.

These considerations of selection on instability and non-symmetrical mutations results in our first-order approximation of the evolution of instability for a minimal cancer cell population:

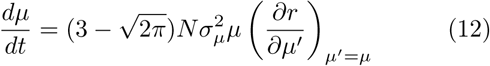

which defines the canonical equation for unstable cancer dynamics under the constraints considered here.

## V. EVOLUTION OF INSTABILITY

The canonical equation (12) describes the evolution of instability in our model population depending on the population size *N*, the distribution of mutation jumps 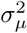 and the product of the mutation rate and the gradient of the adaptive landscape *μ∂_μ_r.* For our model landscape (eq. 5), this turns out to be

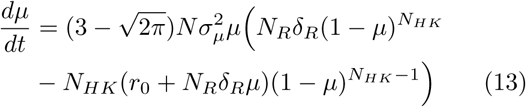

Complicated analytical solutions for this equation might not give best insight of the underlying dynamics. However, as a first test of our model we compare its numerical solution to averaged Moran Process simulations (Fig. 3). It is both relevant and useful to understand the factors that cause deviations between computer experiments and our analytical approach, in order to further comprehend the approximations on which AD is build.

**FIG. 3:**
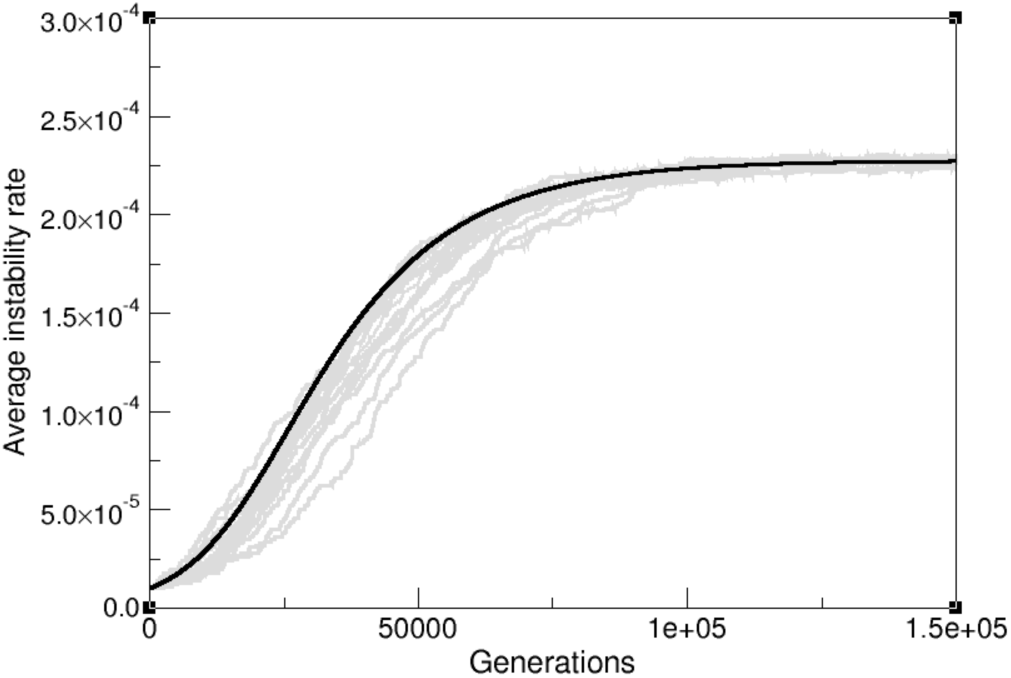
Evolutionary trajectories of the simulated Moran process (gray lines) and numerical solution of the predicted Adaptive Dynamics result (black curve).

These are mostly translated into the population being large enough, and mutation rates being proportionally small. The second is easily met for both healthy and cancerous human cells, but simulating full-size clinically detectable tumours (more than 10^8^ cells (Bozic et al., 2013)) is of large computational cost, and keeping our model and exercise minimal we have used smaller populations, modeling smaller subclones or spatially segregated populations where drift comes into play. Such drift produces evolution to deviate from the gradient trajectory and so proceed slightly slower than our estimate. Together with this, the high nonlinearity of our landscape ensures that *(a_1_(µ)> = a_1_(〈µ〉)* will be only valid up to a certain degree of approximation.

A better understanding of the underlying dynamics can result from dividing the exploration of the landscape in well-behaved regions where simpler equations will arise. On the one hand, in an initial phase of malignancy exploration for small values of μ, the shape of the adaptive landscape is dominated by the linear increase of mutated oncogenes, r(µ) = r_0_ + *N_R_δ_R_µ.* Within this region, dynamics of instability follow

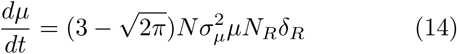

and the mean evolutionary trajectory is

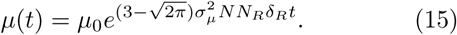

It is remarkable to understand how, even in a linear adaptive landscape, the coupling between mutation and selection on unstable cells introduces a further nonlinearity that will account for exponential exploration of the space of instability and the consequent exponential increases in replication capacity. Such results can be again compared to computer simulations of mutator-replicator cells (Fig. 4). The smaller nonlinearity also ensures that AD remains a good approximation despite stochastic deviation.

**FIG. 4:**
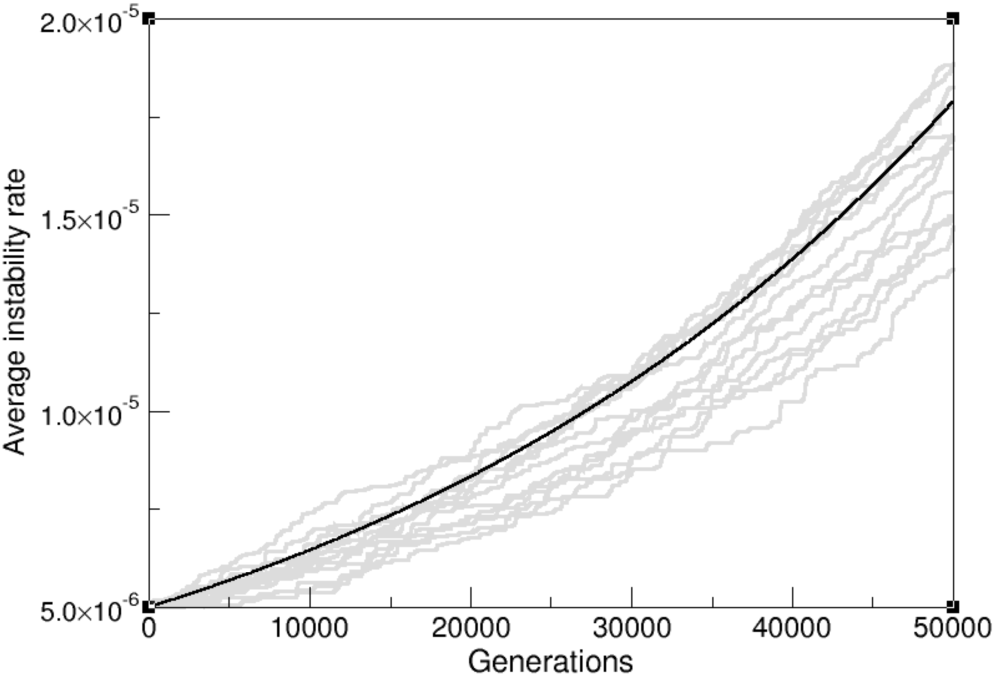
Exponential evolution of the mean mutation rate on a linear landscape: Moran process simulations (gray lines) of populations of 2000 cells and the AD approximation (black curve).

Another interesting point is to understand the behaviour of the mean instability levels as the population approaches the landscape peak. This kind of behaviour is easily studied if one considers a simple landscape containing a peak, such as *r(y) = r_0_ + δ_R_N_R_μ-δ_HK_N_HK_μ^2^*, where the role of house-keeping genes is not considered totally deleterious but just reducing fitness quadratically with the mutation rate. This landscape has an optimal value at p* *= SRNR/2S_HK_*N_hk_, and this peak is explored through

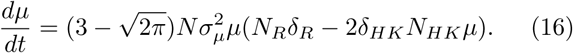

By means of rewriting this trajectory as dμ/dt *= Aμ(B —* Cμ), with 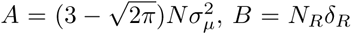 and *C = 2δ_HK_N_HK_*, its solution simplifies to

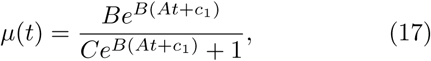

where *c*_**1**_ ensures that µ(0) = µ_0_, the normal mutational rate of healthy cells. This trajectory saturates for long times at the expected result A/B = µ*, and can be again compared to computational experiments of replicating cells (Fig. 5).

**FIG. 5:**
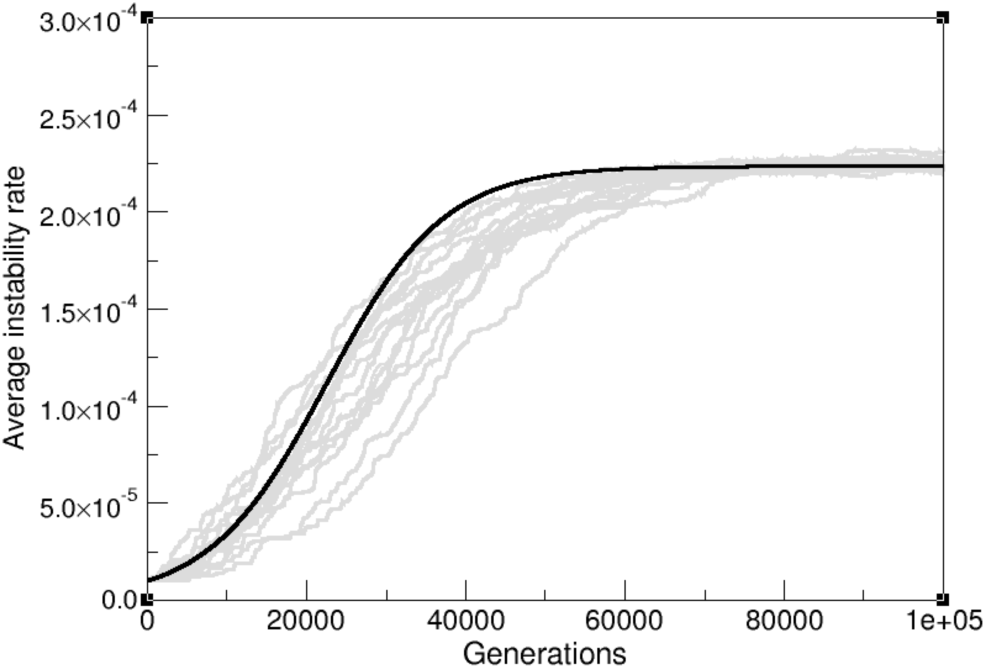
Mean mutation rate saturation at the fitness peak: Moran process simulations (gray lines) of populations of 2000 cells and the AD approximation (black curve).

The same deviation between simulations and the numerical fit is found in this case, with evolution proceeding slower than our estimate. However, this minimal landscape approximation is able to capture the dynamical behaviour of our gene-related landscape model, mainly with an initial exponential growth followed by saturation around the peak, which can be proven to be an evolutionary stable strategy (Geritz et al., 1998).

Provided that the canonical equation has a non-trivial, singular point, as we found for *µ\** = δ_R_N_R_/2S_HK_N_HK_, one can study the evolutionary stability of a quantitative trait. We can easily compute if this singular mutation rate will be an evolutionary trap, *i.e,* a strategy that no further mutants can invade, if

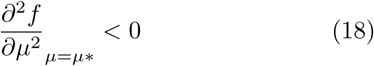

which holds for our strategy: ∂_μμ_r(p) = –2δ_HK_N_HK_ < 0.

## VI. DISCUSSION

In the present work we have discussed the implications of the coupling between selection and instability for a minimal replicator-mutator model. We have shown how to determine the evolutionary trajectory for the mean instability levels in a basic landscape of cancer-related genes. Our AD model, as defined by our canonical equation (and consistently with simulated trajectories) describes the tempo and mode at which mutation rates increase and saturate around fitness peaks. For a simple but sensible fitness landscape, a general canonical equation has been derived from the Moran process scenario. Several approximations have also been considered.

A first relevant result of our model arises from evaluating the canonical equation for unstable cells in a linear landscape, to be associated with a pre-malignant stage. The nonlinearity resulting from the coupling of mutation and selection predicts an exponential increase of instability levels, whereas a trait different from instability would only increase linearly within such landscape. This result is presented as a mathematical description of genomic instability being an enabling characteristic of cancer, by means of generating fast exploration of the space of possible mutations towards malignancy. Similarly, we obtained consistent matchings between simulated and average predicted instability values for the near-optimum state. The possible applications of such minimal evolutionary descriptions of tumour instability follow from our set of examples and computer simulations.

Mounting evidence indicates that a successful approach to cancer therapy requires an explicit evolutionary perspective (Gatenby et al., 2009). One possible instance of this is provided by mutagenic therapies, that have produced key results in the field of virology (Loeb et al., 1999). Would they be effective for cancer? Given some key analogies between RNA virus populations and unstable tumors (Solé et al., 2004) this is an appealing possibility, although drug design or resistance mechanisms have yet to be assessed (Fox et al., 2010). Prior to that, conceptual questions arise, such as: do cancer cells live near critical instability levels, beyond which viability is no longer possible? is there a sharp error threshold for the mutation rate? what evolutionary outcomes should we expect when inducing variations on the mutational load of cancer cells, and how can these shed new light on mutagenic therapy?

Regarding the later, our model includes a novel path to incorporate instability as the evolving trait, while providing potential insights, particularly before and beyond the optimal instability levels, where evolution of instability happens at different rates. The exponentially fast increase of small mutational loads indicate that reducing instability rates in hope for progression delay might result in rapid reexploration of the mutator phenotype. On the other hand, pushing instability beyond optimal levels, even if a critical point is not trespassed (Solé et al., 2004), might render tumor cells too unstable and, what is more, evolution towards more stable regions by means of recovering DNA maintenance mechanisms might prove impossible.

Together with specific mutational-oriented treatment, the importance of instability on both disease prognosis and evolutionary dynamics towards malignancy asks for dynamical descriptions like the one here discussed. Conceptual understanding of cancer adaptation can gain insight by means of considering that mutation and selection are coupled, and prediction of therapy outcome must take into account the increased pace at which cancer cells explore the pathways towards resistance.

While searching for the minimal ingredients needed to build an evolutionary description of unstable tumour populations we have left aside many relevant considerations. This model, therefore, coarse-grains tumour ecology to simple mutator-replicator populations, and produces solely a description of the mean mutation rate evolution. Cancer is more complex, with spatial heterogeneity being a main hallmark of tumour architecture. AD frameworks could better describe heterogeneity by means of considering tumour ecology as a set of different populations, spatially distributed, deploying different hallmark capabilities and coevolving through different instability schemes. A minimal mean-field approach to these sub-clonal dynamics, like the one here presented, can be a first step towards better understanding and predicting unstable intratumoural coevolution.

## Footnotes

1 The Rayleigh distribution is an asymmetric probability distribution defined for positive random variables (Forbes et al., 2010). We displaced so that it is Δμ = 0 peaked, with shape 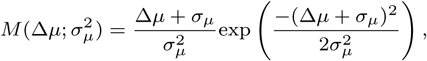 accounting for asymmetric probabilities of possible mutation levels, with small forward mutations being the most common.

## ACKNOWLEDGMENTS

The authors thank P. Ruiz and E. Beltran, as well as the CSL lab members for useful discussions. Special thanks to Oliver Law for his inspiring ideas. This work has been supported by the Botín Foundation by Banco Santander through its Santander Universities Global Di-vision, a MINECO grant FIS2015-67616 fellowship, by the Universities and Research Secretariat of the Ministry of Business and Knowledge of the Generalitat de Catalunya and the European Social Fund, and by the Santa Fe Institute.

